# Diversity, dynamics and effects of LTR retrotransposons in the model grass *Brachypodium distachyon*

**DOI:** 10.1101/710657

**Authors:** C Stritt, M Wyler, EL Gimmi, M Pippel, AC Roulin

## Abstract

- Transposable elements (TEs) are the main reason for the high plasticity of plant genomes, where they occur as communities of diverse evolutionary lineages. Because research has typically focused on single abundant families or summarized TEs at a coarse taxonomic level, our knowledge about how these lineages differ in their effects on genome evolution is still rudimentary.
- Here we investigate the community composition and dynamics of 32 long terminal repeat retrotransposon (LTR-RT) families in the 272 Mb genome of the Mediterranean grass *Brachypodium distachyon.*
- We find that much of the recent transpositional activity in the *B. distachyon* genome is due to centromeric *Gypsy* families and *Copia* elements belonging to the Angela lineage. With a half-life as low as 66 ky, the latter are the most dynamic part of the genome and an important source of within-species polymorphisms. Second, GC-rich *Gypsy* elements of the Retand lineage are the most abundant TEs in the genome. Their presence explains more than 20 percent of the genome-wide variation in GC content and is associated to higher methylation levels.
- Our study shows how individual TE lineages change the genetic and epigenetic constitution of the host beyond simple changes in genome size.

## Introduction

Transposable elements (TEs) are stretches of DNA which can replicate within genomes (Burt & Trivers, 2006). Since their discovery in the 1950ies by Barbara McClintock, their fuzzy status between selfish parasite and integral part of the host genome has puzzled biologists. In contrast to viruses, TEs do not routinely leave the host; instead, their evolutionary history is largely one of vertical transmission and co-evolution in a host genome which has evolved epigenetic mechanisms to suppress their activity (Lisch, 2009). As suggested by the omnipresence of TEs in the eukaryote domain, this co-evolution is ancient and has shaped eukaryote genome evolution from the very beginning (Eickbush & Malik, 2002).

Over the past two decades, comparative and functional genomics studies have revealed how TEs can inflate genome size (Ma & Bennetzen, 2004; Piegu *et al.*, 2006) and produce novel host phenotypes by inserting into genic regions (van’t Hof *et al.*, 2016; Niu *et al.*, 2019). While these examples have added to the sense that TEs play an important role in evolution (Casacuberta & González, 2013; Belyayev, 2014), it has also become clear that TEs are immensely diverse, and that what is found in one species does not necessarily hold in others. This is particularly true for plants, whose genomes are subject to fewer evolutionary constraints than those of animals (Kejnovsky *et al.*, 2009) and harbour diverse TE lineages with different structures and replication strategies (Wicker & Keller, 2007; Du *et al.*, 2010; Neumann *et al.*, 2019). TE landscapes can diverge rapidly because TE activity depends on multiple interacting factors whose direction and strength can differ within a single species. These factors include environmental triggers (Horváth *et al.*, 2017), the demographic history of the host population (Lynch, 2007), and horizontal transfers (Baidouri *et al.*, 2014).

Such complexity and historical contingency is the hallmark of ecological systems, and it has therefore been proposed to borrow concepts from ecology and conceive of genomes as ecosystems inhabited by various ‘species’ of TEs differing in behaviour and genomic niches (Kidwell & Lisch, 2001). While the scope of TE ecology is not very clear-cut (Brookfield, 2005; Venner *et al.*, 2009; Linquist *et al.*, 2013), one important intuition this metaphor conveys is that our understanding of genome evolution can be improved by taking the diversity of TEs into account rather than lumping them together into coarse taxonomic units (Stitzer *et al.*, 2019). Most studies with an interest in mobile elements consider TEs at the class (DNA transposons vs. retrotransposons) or superfamily (*Copia* vs. *Gypsy*) level. Such comparisons provide valuable overviews of TE communities and have revealed some general patterns, e.g. that DNA transposons tend to occur closer to genes than retrotransposons (Feschotte & Pritham, 2007). Yet, considering that for example the *Gypsy* superfamily consists of more than a dozen of lineages which may be as old as the major divisions of plants (Neumann *et al.*, 2019), generalisations about ‘repeats’, ‘retrotransposons’, or ‘*Gypsy* elements’ are liable to level out biologically important differences between TE lineages.

With the increasing amount of information available for some model organisms, it has become possible to investigate a single genome ecosystem in detail by relating TEs to recombination rate, methylation levels, and other properties of the genomic context (Stitzer *et al.*, 2019). In this study we use the excellent genomic resources available for the wild Mediterranean grass *Brachypodium distachyon* in order to investigate the long terminal repeat retrotransposons (LTR-RT) in this species. LTR-RTs, the most abundant TEs in plants, are typically between 5 and 15 kb long and multiply through a copy-and-paste mechanism involving an RNA intermediate (Eickbush & Malik, 2002). Apart from the reverse transcription step, which LTR-RTs share with retroviruses, the two LTRs flanking these insertions are the most characteristic feature of this class of TEs. These sequences do not only allow a comparatively easy identification of these repeats in the genome, but also play a vital role in LTR-RT life history as they contain regulatory motifs (Schulman, 2013) and are prone to ectopic recombination. In *B. distachyon*, LTR-RTs make up 20 percent of the 272 Mb genome and are widely dispersed along the five chromosomes (International Brachypodium Initiative, 2010; Schulman, 2015). This modest amount of repeats, compared to the TE ‘jungles’ of wheat, maize, and other large-genome plants, allows studying the LTR-RT community without loosing sight of its constituent ‘species’.

The goal of our study is to provide an overview of the LTR-RT community in *B. distachyon* and to characterize its dominant lineages and their effect on genome composition and dynamics. In particular, we address the following questions. Which major plant LTR-RT lineages are present in *B. distachyon*, and what is their relative abundance and age? How are these different lineages related to important genomic features such as genes, recombination rate, methylation, GC content, and genetic diversity?

## Material and Methods

### Genome assemblies

Two genome assemblies for *Brachypodium distachyon* are considered in this study. While the focus is on the reference accession Bd21 for which most information is available, a new assembly for the Turkish accession BdTR7a is included in order to investigate within-species differences in LTR-RT communities. The assembly for the reference accession Bd21 (version 3.0) was downloaded from Phytozome 12; it is based on BAC libraries and has well-assembled repetitive regions (International Brachypodium Initiative, 2010; VanBuren & Mockler, 2016). We chose BdTR7a to create a second assembly because among the 54 recently sequenced accessions it has the highest number of non-reference transposable element insertions (Stritt *et al.*, 2018). The BdTR7a assembly was created by combining PacBio sequencing with Bionano optical mapping (Supporting Information Methods S1).

### Annotation of LTR retrotransposons and reverse transcriptase phylogeny

Because a consensus library for the different TE families of *B. distachyon* is available on the TREP database (http://botserv2.uzh.ch/kelldata/trep-db/index.html), we used these sequences as a starting point to annotate LTR-RTs. For the sake of consistency, we used the same approach to annotate TEs in the new assembly and to re-annotate them in the reference genome. The LTR sequence of each of the 21 *Copia* and 19 *Gypsy* consensus sequences was blasted against the assemblies. Hits which covered at least 80 percent of the LTR were retained and sorted according to their position on the chromosome. We then traversed the sorted hits and compared adjacent LTR pairs. A hit pair was denoted *intact* if the two hits belonged to the same family, were on the same strand, and the distance between them corresponded to the distance expected from the consensus sequence, with an error margin of 20 percent to account for indels. Otherwise the hit was denoted a *single* LTR. A single LTR was classified as *solo* LTR if it was flanked by identical 4-mers, being evidence for a target site duplication (TSD), and lacked internal TE sequence in its 500 bp flanking regions. For the comparison of intact and solo elements, we only included intact elements satisfying the same stringent criteria, in this case requiring TSDs and the presence of internal TE sequence 500 bp up- or downstream of the LTRs. The Python script implementing this annotation method is available on github.com/cstritt/tes.

In order to determine the evolutionary relatedness of the annotated TE families and their place in the larger phylogeny of plant LTR retrotransposons, we searched the six-frame translated TE consensus sequences against the Pfam database (pfam.xfam.org) with the HMMER tool hmmscan (Finn *et al.*, 2011) and extracted the sequences aligning to the reverse transcriptase profiles RVT_1 (*Gypsy*) and RVT_2 (*Copia*). Amino acid sequences of the major *Copia* and *Gypsy* lineages in plants were obtained from the RepeatExplorer database (Neumann *et al.*, 2019). A reverse transcriptase consensus sequence for each lineage was constructed after aligning the individual RT copies with MAFFT v.7.402 (Katoh & Standley, 2013). *Copia* and *Gypsy* consensus sequences were then merged with the respective *B. distachyon* RT sequences, aligned with MAFFT (--auto), and trees were estimated with MrBayes 3.2.2 (Ronquist *et al.*, 2012) by sampling over different amino acid models (aamodelpr = mixed) and running two chains for 500,000 generations. Tracer (Rambaut *et al.*, 2018) was used to assess the convergence and mixing of the MCMC runs. Trees were visualised with Figtree (tree.bio.ed.ac.uk/software/figtree).

### Estimation of insertion age and family survival functions

To estimate the age of intact and solo insertions, we used the LTR sequences to construct LTR genealogies. LTRs from single and intact elements for each family were aligned with MAFFT (-- auto) and trimmed with trimal (Capella-Gutiérrez *et al.*, 2009) to remove sites with more than 5% gaps in the alignment. Trees were estimated with the uniform clock model in MrBayes, with a HKY substitution model (nst=2) and an inverse gamma prior on the clock rate (rates=invgamma). From the LTR trees we extracted the terminal branch lengths as a proxy of insertion age. In the case of intact elements, this corresponds to well known method of age estimation from LTR divergence (SanMiguel *et al.*, 1998); for solo LTRs, the terminal branch length represents an upper-limit age estimate as it represents the time to the most recent common ancestor of the solo LTR and another copy rather than the two solo LTRs of a single copy.

Assuming that full-length copies die at a constant rate, the survival function of a LTR-RT family can be approximated by an exponential function of the form N_t_ = N_0_ e^− λt^, where λ is the death rate of the family. To estimate λ for the nine families with at least 20 full length copies, exponential distributions were fitted to the age distributions using the *fitdistr* function of the R package MASS, and the exponential rate was recovered. In addition, we estimated λ_S_, an alternative death rate estimate which includes information on the age of solo LTRs, using the maximum likelihood function of Dai et al. (2018). Confidence intervals for λ_S_ were obtained from 100 bootstraps. Both procedures were implemented in R and are available on github.com/cstritt/tes.

### Genomic niche features

In order to characterize the genomic niches of LTR-RT lineages we compiled a data set of diverse genomic features which might affect or be affected by the presence of LTR-RTs. Recombination rates were obtained from the linkage map of Huo et al. (2011). Distance to the closest gene was calculated based on v.3.1 of the reference genome annotation, downloaded from Phytozome 12. Copy methylation levels were obtained through whole-genome bisulfite sequencing (Supporting Information Methods S2). Finally, we estimated three population genetic statistics in 10 kb windows around the annotated TEs, 5 kb on each side: Kelly’s Z_nS_, a measure of multilocus linkage disequilibrium and thus a proxy for local recombination rates; Tajima’s D, a statistic indicating deviations from neutrality; and the number of segregating sites S and nucleotide diversity π. Variants for 25 Turkish and Iraqi accessions were extracted from the Phytozome 12 variant set, and the R package *PopGenome* (Pfeifer et al., 2014) was used to estimate the statistics.

### Statistical inference

To test whether the ratio of solo- to full-length elements can be predicted by the length of the LTR and the internal TE sequence, we fitted a generalised linear model with a binomial error distribution, using the *glm* function in R. Since TE families are phylogenetically related and not independent observations, we used only one randomly chosen family per lineage, resulting in a total of 12 observations. The Tekay family RLG_BdisC004 was excluded because of its unusually long LTR.

A random forest approach (Breiman, 2001) was used to discover features of the genomic context associated to the occurrence of specific TE lineages. We preferred this method over a parametric modelling approach because model specification proved impracticable given the high level of variable collinearity, phylogenetic signal, and chromosomal autocorrelation in our genome-wide data set. 200 decision trees were grown using the *RandomForest* package in R (Liaw & Wiener, 2002) to predict which TE lineage is present at an insertion site as a function of the genomic niche features described above. Predictor variables were ranked according to the mean decrease in accuracy, a statistic which describes how much worse the model performs when the values of each variable are permuted.

To investigate the relationship between TE copy age and GC content, a linear mixed model with the TE lineage as random effect was fitted with the *lmer* function of the R package lme4. All statistical analyses were done with R v3.4.4.

### TE copy synteny and polymorphisms

To test whether TE dynamics differ within the species, we compared annotated TEs in Bd21 and BdTR7a, two eastern Mediterranean accessions whose ancestors diverged between 0.5 and 1.5 mya (Sancho *et al.*, 2017). The discordant read-pair approach implemented in *detettore* (http://github/cstritt/detettore) was used to test whether a copy annotated in one accession is present in the other accession or not. Illumina paired-end reads for the two accessions were downloaded from the NCBI database (SRA samples SRS1615350 and SRS360854) and aligned to the two respective genome assemblies with BWA MEM (Li, 2013). *Detettore* was then used to detect clusters of read pairs with abnormal insert sizes which span annotated TEs. A table containing annotated TEs, the values for their genomic niche features, as well as their conservation state between Bd21 and BdTR7a is provided in the supplement (Tables S1 & S2).

## Results

### Major lineages of LTR retrotransposons in *B. distachyon*

Dotplots and pairwise alignments of the 40 LTR retrotransposon consensus sequences from the TREP database revealed the presence of elements with sequence similarities higher than 80 percent and thus not classifying as families *sensu* Wicker et al. (2007, Fig. S1). Such elements were merged and given the name of the most abundant subfamily, resulting in a total of 32 families (17 *Copia* and 15 *Gypsy*) which were analysed in this study.

Phylogenetic trees based on the reverse transcriptase (RT) sequences show that eleven major lineages of LTR-RTs are present in *Brachypodium distachyon* (Fig. 1). Some lineages are present as a single family, namely the *Copia* lineages Bianca, SIRE, Alesia, and Ikeros, as well as the *Gypsy* lineages Reina and Tekay; others comprise multiple families, notably the six *Copia* families of the Ivana lineage and the three families of the Angela lineage, as well as the nine *Gypsy* families of the Retand lineage and the three families of the CRM lineage (Fig. 1). The Retand families are further divided into three subclades of which subclade C, with four families and 139 full length elements, is the most abundant and diverse. Interestingly, the *Gypsy* element with most full length copies, RLG_BdisC152, could not be classified based on reverse transcriptase as no sequence homology was found. Indeed, apart from a *gag* fragment, this family lacks retrotransposon domains altogether. A hint to its origin comes from a stretch of 2000 bp including the 3’ half of the LTR and the adjacent region, which has high similarity to the CRM families (Fig. S2).

**Fig. 1:**
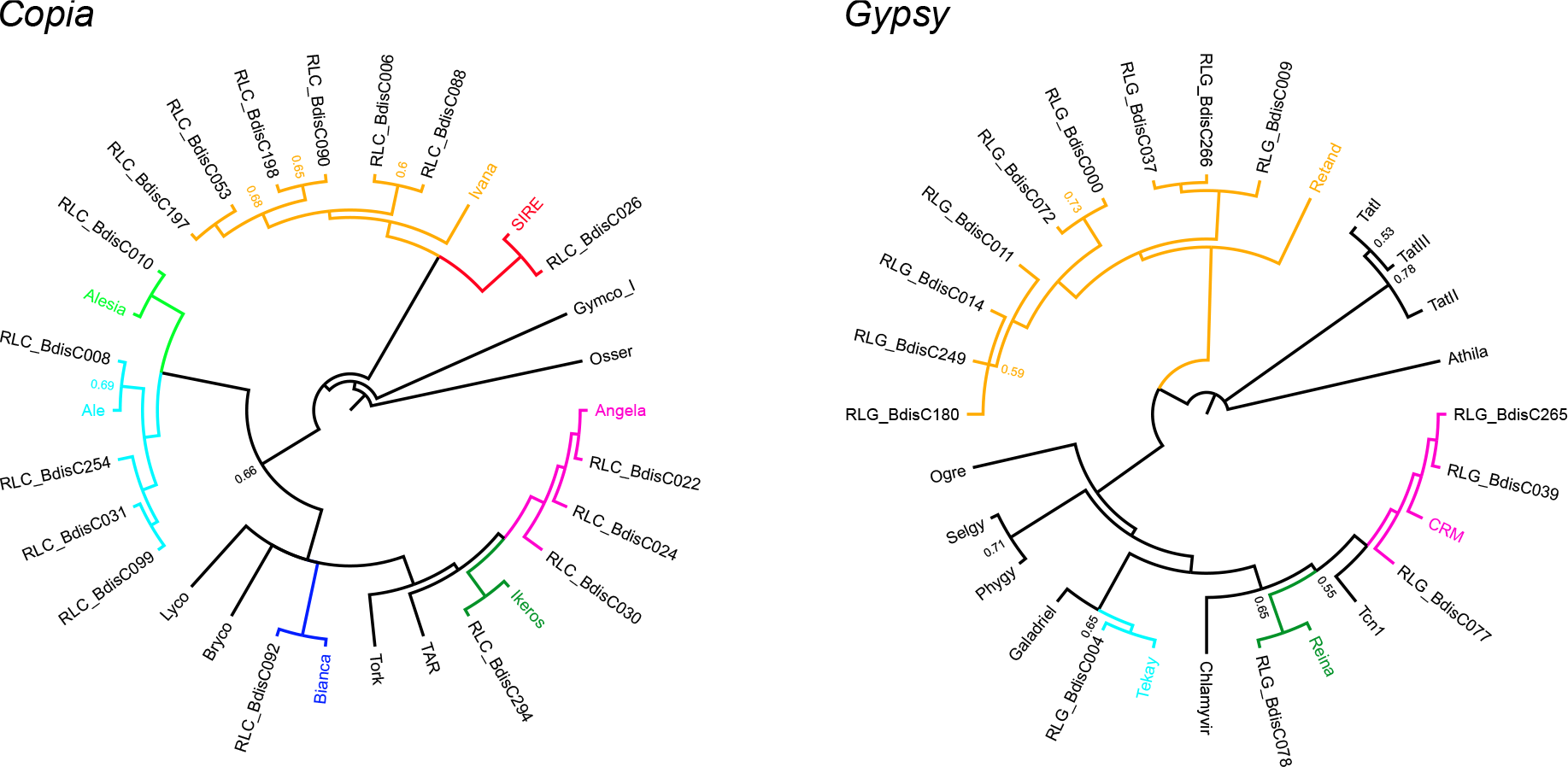
Phylogeny of LTR retrotransposons based on reverse transcriptase sequences. Tip labels not beginning with RL represent consensus sequences of major plant TE lineages; the coloured clades show the lineages present in *B. distachyon*. Posterior probabilities for splits are only indicated when they are lower than 0.8. Absent from this figure is RLG_BdisC152, which lacks a reverse transcriptase domain.

### Community composition

The abundance of the 32 LTR-RT families in the reference accession Bd21 varies from 0 (RLG_BdisC037) to 58 full length elements (RLG_BdisC152, Table 1, Fig. 2). The three most abundant *Copia* families, all of which belong to the Angela lineage, have between 27 and 45 full length elements. The Alesia family RLC_BdisC010 is similarly abundant (24 intact copies), while most other *Copia* families are present in low copy numbers (<10). *Gypsy* elements are generally more abundant than *Copia* elements, the largest contribution coming from families of the Retand lineage, the CRM family RLG_BdisC039, and the unclassified RLG_BdisC152.

**Table 1.** Overview of annotated LTR-RT families in the reference accession Bd21.

| Lineage | TE family | Length | LTR length | GC | Intact<sup>1</sup> | Solo | Unclassified | S/I<sup>2</sup> |
| --- | --- | --- | --- | --- | --- | --- | --- | --- |
| *Copia* |  |  |  |  |  |  |  |  |
| Angela | RLC_BdisC024 | 7,917 | 1,355 | 47 | 27 \| 20 | 160 | 71 | 8 |
|  | RLC_BdisC022 | 8,676 | 1,712 | 44 | 45 \| 32 | 95 | 46 | 2.97 |
|  | RLC_BdisC030 | 7,955 | 1,328 | 47 | 28 \| 22 | 176 | 133 | 8 |
| Ikeros | RLC_BdisC294 | 6,219 | 420 | 45 | 7 \| 0 | 3 | 65 | NA |
| Bianca | RLC_BdisC092 | 5,254 | 228 | 38 | 3 \| 3 | 1 | 15 | 0.33 |
| Ivana | RLC_BdisC197 | 5,239 | 375 | 49 | 6 \| 6 | 0 | 1 | 0 |
|  | RLC_BdisC053 | 5,350 | 409 | 50 | 8 \| 8 | 1 | 5 | 0.13 |
|  | RLC_BdisC198 | 5,210 | 417 | 53 | 4 \| 4 | 0 | 0 | 0 |
|  | RLC_BdisC090 | 5,200 | 366 | 60 | 6 \| 5 | 0 | 0 | 0 |
|  | RLC_BdisC006 | 5,407 | 498 | 50 | 2 \| 2 | 0 | 2 | 0 |
|  | RLC_BdisC088 | 5,161 | 338 | 54 | 4 \| 3 | 1 | 1 | 0.33 |
| SIRE | RLC_BdisC026 | 9,276 | 1,345 | 39 | 10 \| 9 | 131 | 138 | 14.56 |
| Ale | RLC_BdisC099 | 4,796 | 200 | 48 | 4 \| 3 | 0 | 3 | 0 |
|  | RLC_BdisC031 | 4,846 | 223 | 50 | 4 \| 4 | 0 | 2 | 0 |
|  | RLC_BdisC254 | 4,915 | 208 | 57 | 3 \| 3 | 0 | 3 | 0 |
|  | RLC_BdisC008 | 5,170 | 132 | 52 | 5 \| 4 | 2 | 7 | 0.5 |
| Alesia | RLC_BdisC010 | 5,894 | 720 | 45 | 24 \| 16 | 30 | 10 | 1.88 |
| *Gypsy* |  |  |  |  |  |  |  |  |
| Retand A | RLG_BdisC009 | 13,537 | 821 | 59 | 22 \| 9 | 31 | 87 | 3.44 |
|  | RLG_BdisC266 | 13,197 | 915 | 56 | 45 \| 26 | 116 | 230 | 4.46 |
|  | RLG_BdisC037 | 15,485 | 854 | 55 | 0 \| 0 | 2 | 2 | NA |
| Retand B | RLG_BdisC000 | 13,500 | 808 | 55 | 57 \| 31 | 81 | 349 | 2.61 |
|  | RLG_BdisC072 | 14,246 | 478 | 53 | 10 \| 5 | 7 | 69 | 1.4 |
| Retand C | RLG_BdisC011 | 14,104 | 972 | 66 | 47 \| 29 | 75 | 316 | 2.59 |
|  | RLG_BdisC014 | 12,887 | 838 | 63 | 26 \| 14 | 41 | 260 | 2.93 |
|  | RLG_BdisC249 | 13,155 | 915 | 62 | 17 \| 4 | 37 | 203 | 9.25 |

**Fig. 2:**
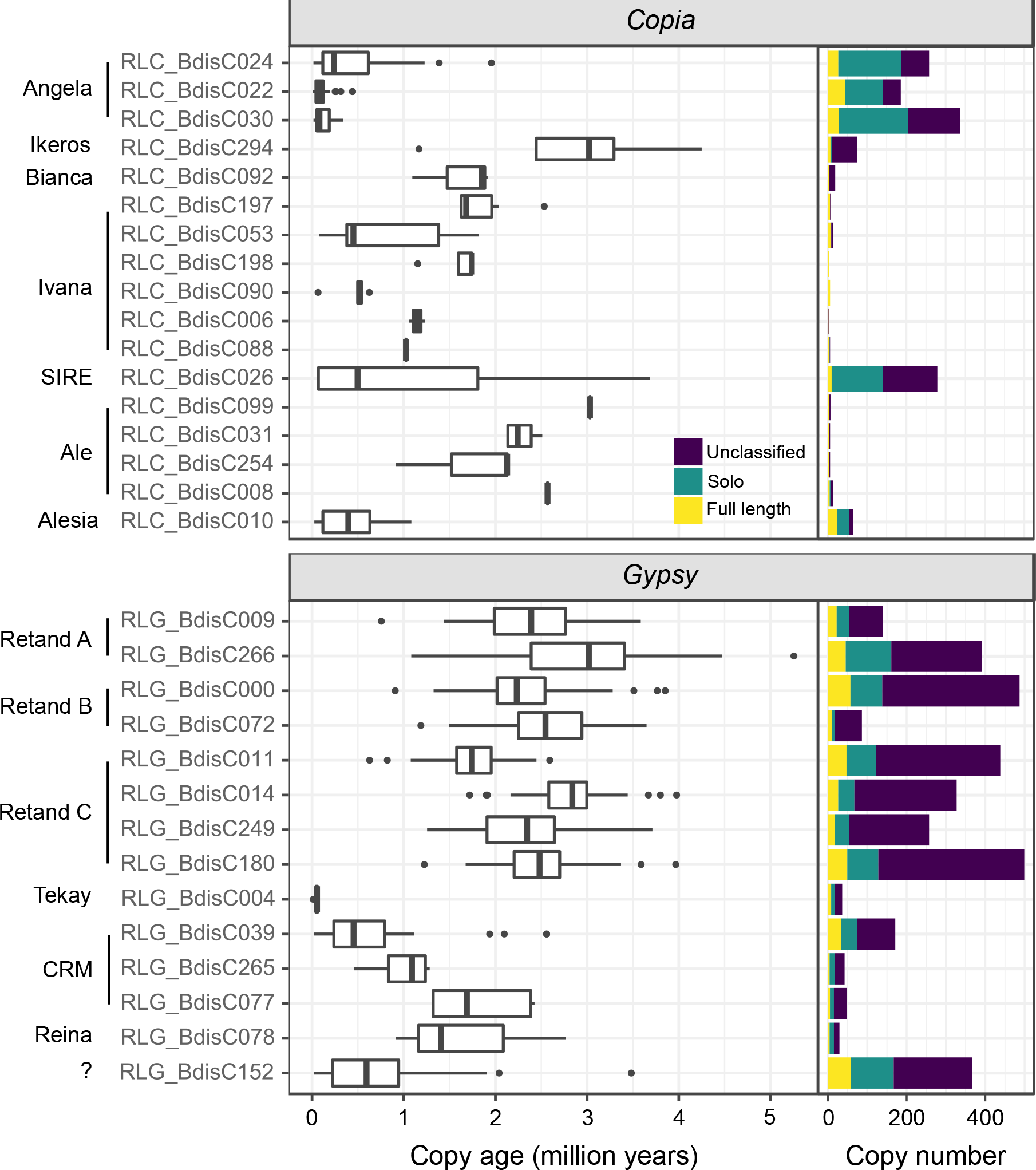
Age distributions of full-length elements and copy numbers of LTR-RT families. *Copia* lineages and families in the upper panel, *Gypsy* in the lower. The Retand family RLG_BdisC037 is missing on this figure because it has no full length copies.

In most LTR-RT families, the number of solo LTRs exceeds the number of intact elements (Table 1, Fig. 2). An exception are several low copy-number *Copia* families for which we detected only few or no solo LTRs. High solo-to-intact (S/I) ratios are associated to long LTRs (r = 0.73, P<0.001) and high ratios of LTR versus internal sequence length (r = 0.47, P=0.01). The significance of the latter association is supported by a binomial generalized linear model which includes LTR and internal sequence length as well as their interaction term as predictor variables: while LTR length itself is not significant in this model (P=0.55), the interaction term is (P=0.04), indicating that high rates of solo LTR formation are favoured by a combination of long LTRs and short internal sequences separating them.

Finally, a considerable number of annotated LTRs could not be classified as being part of a full length or a solo element (Table 1). Annotated Retand LTRs frequently lack target site duplications and/or the expected sequence context that would reveal them as solo LTRs. Among the *Copia* families, a majority of the elements still have these signatures, though the number of unclassified elements surpasses 100 for RLG_BdisC030 and RLC_BdisC026. As shown in the next paragraph, the lack of signatures is associated to the age of the families.

### Age structure of the LTR-RT community and family half-lifes

Insertion age estimates for full-length elements based on LTR divergence reveal a multi-layered age structure of the LTR-RT community in *B. distachyon* (Fig. 2). The Angela families are characterized by low LTR divergence (median copy age = 0.4 mya). For many Angela copies, the 95% credibility interval of the age estimates reaches into the present, illustrating the impossibility to tell whether an element jumped yesterday or a few thousand years ago. The abundant Retand elements, on the other hand, are comparatively old, with a median age of 2.4 my, although this varies for the different families. Ivana and CRM, two other multi-family lineages, show a stepwise age pattern with some old and apparantly inactive families and potentially active families with young copies (RLC_BdisC053, RLG_BdisC039).

Not only average ages vary between lineages and families, but also the range of age estimates. Narrow ranges are partly due to polytomies in the LTR genealogies (Fig. S3): the lack of information in highly similar (Angela) or short (RLC_BdisC008, RLC_BdisC088, RLC_BdisC099) LTRs leads to unresolved nodes in the LTR tree and identical branch lengths for multiple copies. Other families comprise copies of very different ages. Most strikingly, the age distribution of the ten intact SIRE copies spans two million years, and judging from the presence of one young insertion with identical LTRs (RLC_BdisC026_Bd4_6010841), the family is still active.

Age distributions inferred from LTR divergence only provide an indirect picture of TE activity because their shape does not only depend on TE insertion rates, but also on how rapidly full length copies disappear from the genome (Dai *et al.*, 2018). Survival curves for the nine TE families with more than 20 intact elements reveal great differences among families (Table 2, Fig. 3). The half lifes of the three abundant Angela families are the lowest and range from 66 ky (CI=53-91) for RLC_BdisC030 to 152 ky (CI=98-201) for RLC_BdisC024. The two centromeric *Gypsy* families RLG_BdisC039 and RLG_BdisC152 are more persistent, with half lifes of 495 ky (CI=407-854) and 341 ky (CI=298-511). The longest living high-copy-number families belong to the Retand lineage with half-life values between 1,210 ky (CI=871-1,323) and 1,648 ky (CI=1,166-2,026).

**Table 2.**
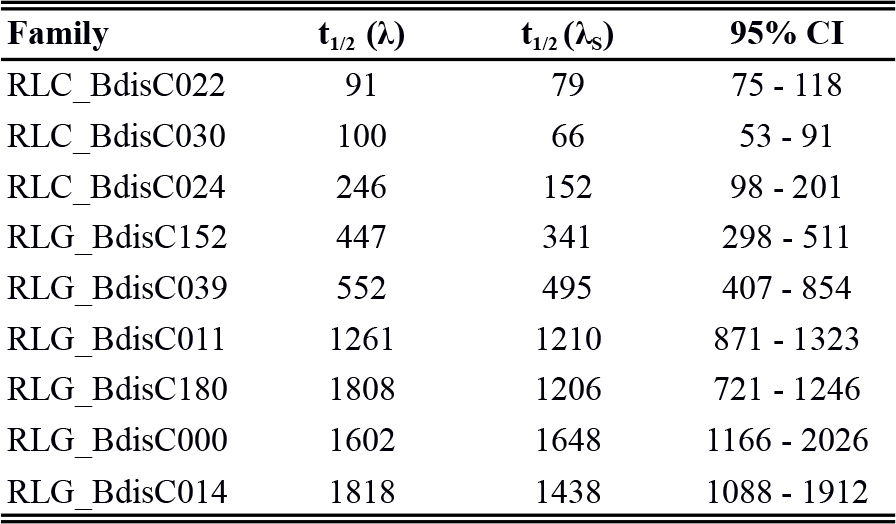
Half-life times for nine high copy-number families. Confidence intervals for the λ_S_ half lifes were obtained from 100 bootstrap replicates.

**Fig. 3.**
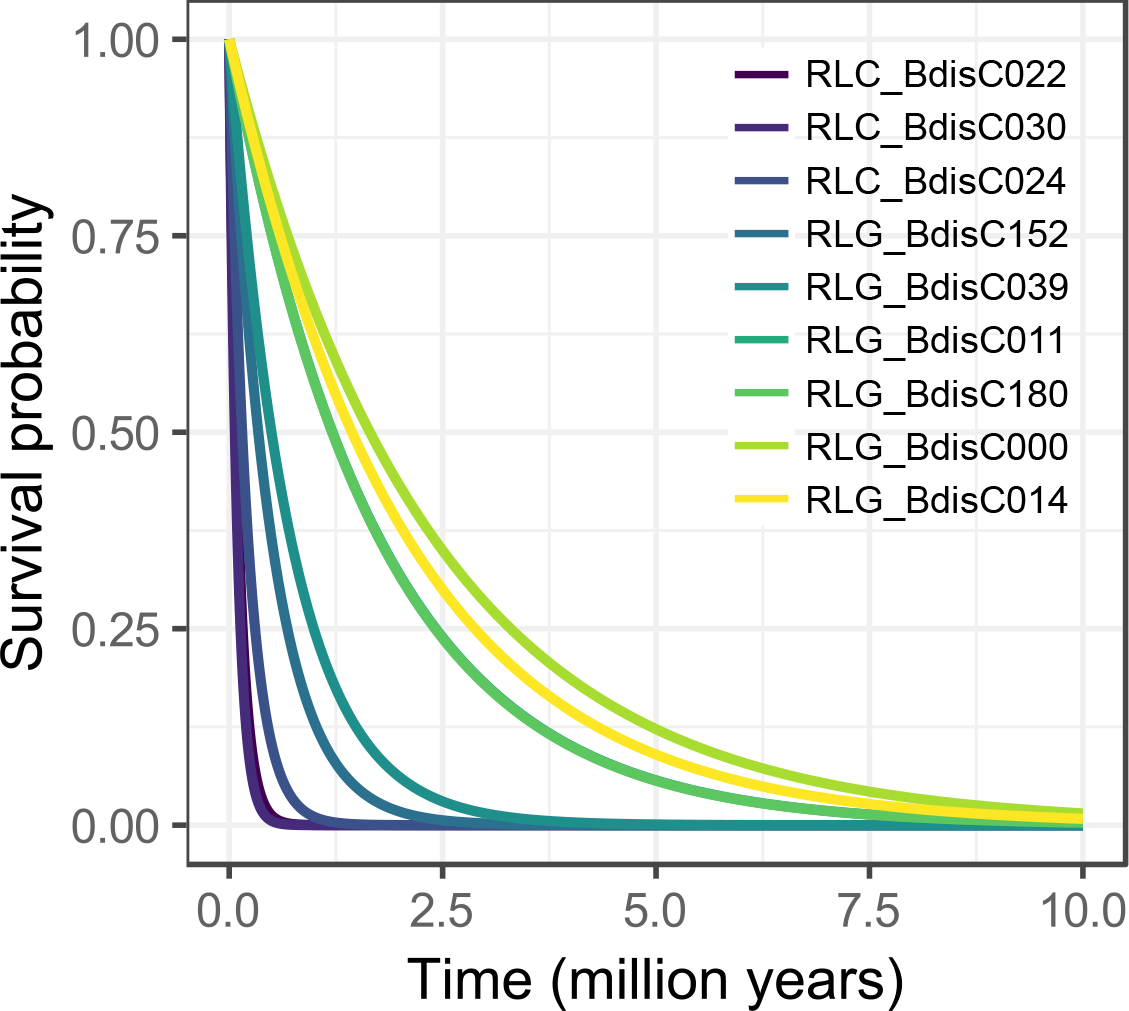
Survival curves of nine high copy-number families.

### TE copy synteny between Bd21 and BdTR7a

In order to find out whether recent TE activity has led to within-species differences in the LTR-RT community, we annotated LTR-RTs in the Turkish accession BdTR7a. The LTR-RT communities of Bd21 and BdTR7a are highly similar, as can be seen in the strong correlation of family abundances between the two accessions (r=0.99, P<0.001, Fig. S4). The largest differences are due to the two Angela families RLC_BdisC022 and RLC_BdisC030, for which 77 and 59 intact elements were annotated in BdTR7a compared to 45 and 26 in Bd21.

456 out of 4612 (9.9%) LTR-RT copies present in Bd21 have no homolog in BdTR7a, 100 of them full length copies, 350 solo LTRs, and 6 unclassified. Strikingly, 235 of these polymorphisms are Angela solo LTRs, compared to 44 intact Angela copies. The other way around, 495 out of 4493 (11%) TEs present in BdTR7a have no homolog in Bd21, 126 of them full length, 357 solo LTRs, and 12 unclassified. Also here more than half of the TE polymorphisms (251) are Angela solo LTRs. Other lineages also contribute to TE polymorphisms: of the annotated Alesia elements, 34 and 42% had no homologs in Bd21 and BdTR7a, respectively. Similar numbers were found for the Tekay family (24 and 42 %), while the CRM lineage (7 and 10%) and the SIRE lineage (13 and 12 %) were more conserved. Almost all annotated elements were conserved between the two accessions in the Ale, Bianca, Ikeros, Reina, and Retand lineages, consistent with their older age (Fig. 2).

Non-conserved elements have a median age of 0.43 my, while for conserved elements this value is 2.5 my. 90% of the non-conserved elements are younger than 1.5 my, and 90% of the conserved elements older than 0.7, which agrees well with a divergence time of the ancestral lineages of the two accessions between 0.5 and 1.5 million years ago (Sancho *et al.*, 2017). The mean distance to genes of the 456 non-conserved elements in Bd21 is 7,897 (SD = 11,694), while for conserved elements the mean is 11,200 (SD = 16,483), a statistically significant difference (Welch’s t-test, P = 6.7e−8).

### Niche characteristics of LTR retrotransposon lineages

The LTR retrotransposons analysed in this study have a non-homogenous distribution along chromosomes (Fig. 4a, Fig. S5). Among the nine lineages with more than 50 annotated elements, three ‘macroecological’ distribution patterns can be distinguished: centromeric (CRM, RLG_BdisC152), centrophobic (Alesia), and centrophilic (Angela, Ikeros, SIRE, Retand). Of these, the close association of the CRM families to the centromere is the strongest pattern and suggests an active targeting mechanism rather than a passive aggregation due to selective constraints. Indeed, in all three CRM families we found the Putative Targeting Domain (PTD) at the integrase C-terminus which is believed to mediate the targeted integration of these elements into the centromere (Fig. S6, Neumann et al., 2011). No such domain was found in the non-autonomous centromere-specific family RLG_BdisC152.

**Fig. 4.**
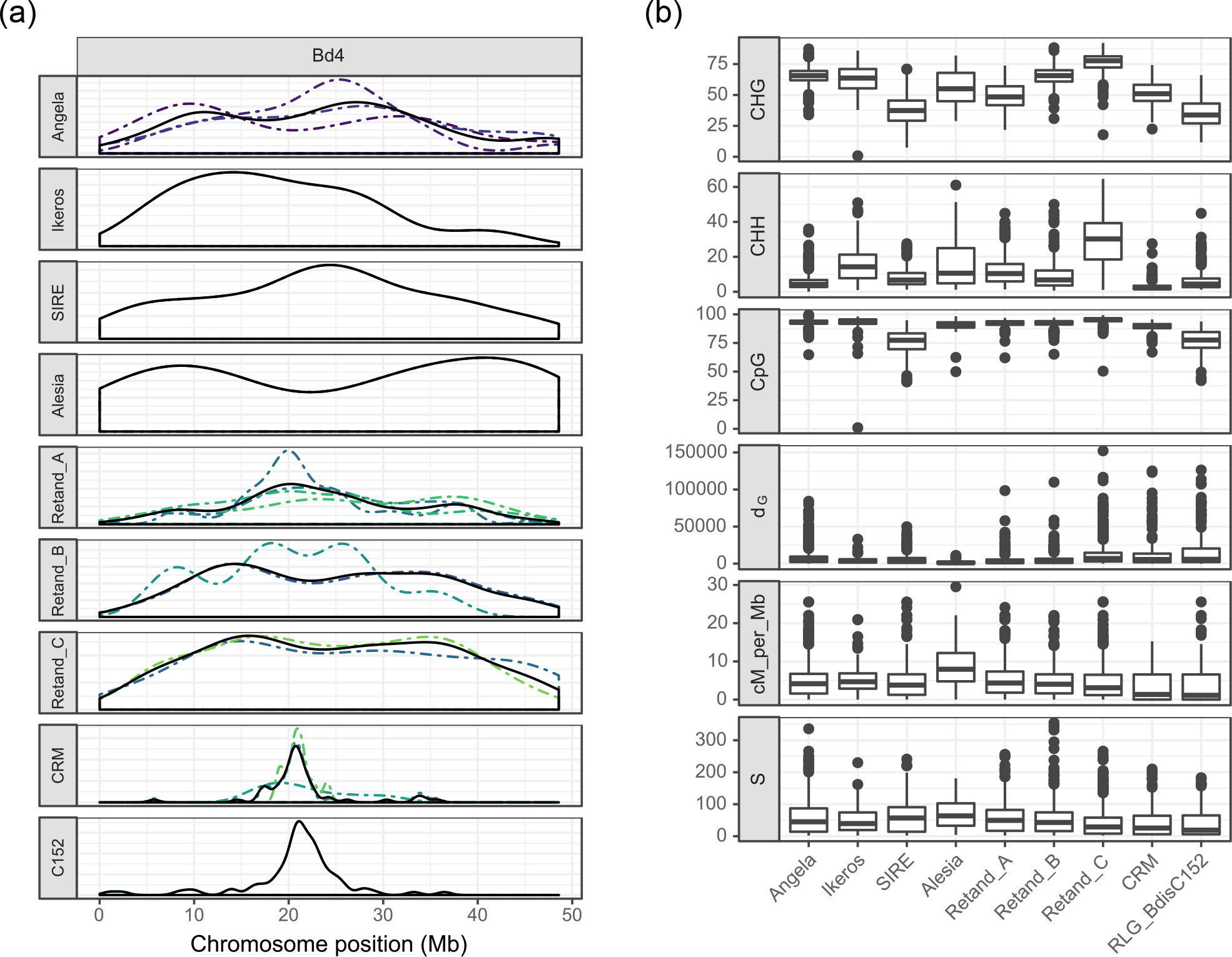
Genomic niches of LTR-RTs. a) Density of the 32 LTR-RT families along chromosome Bd4. Dashed lines indicate the different families within a lineage, the solid line is their average. b) Genomic niche features which best predict the different lineages, ranked according to their importance in the random forest model.

To get a more precise idea about the genomic niches of different TE lineages, we used a random forest model to identify genomic features associated to the occurrence of individual lineages (see Material & Methods). It correctly predicted two thirds of the observations (OOB error rate 31.58%, Tables S3 & S4). The three methylation contexts were by a large margin the best features to distinguish TE lineages, followed by distance to the next gene, recombination rate, and number of segregating sites S (Fig. 4b, Table S4).

Some lineages have copies with a wide range of CHG methylation levels (Fig. 4b), e.g. the Alesia (SD = 14.6) and the Ikeros elements (SD = 12.9); accordingly, they cannot be predicted by their CHG methylation. For other lineages, however, the methylation range is much narrower and quite characteristic: mean CHG methylation is 78% (SD = 7.0) for the Retand clade C and 65% for Angela (SD = 6.7), while it is only 35% (SD = 11.7) for RLG_BdisC152. The same holds for CHH methylation, which correlates with CHG (r=0.57, P<0.001) and distinguishes the same lineages, in particular Retand clade C with 29% (SD = 13.1) compared to an overall mean of 15%. Methylation at CpG sites is generally high (mean 91%) and less variable than the other two contexts. Yet also here two lineages stand out with lower means and variability: RLG_BdisC152 and SIRE with means of 76 (SD = 9.8) and 75 (SD = 10.8), respectively.

Distance to the next gene (d_G_) allows to distinguish the centromere-associated lineages Retand C, CRM and RLG_BdisC152, which have mean d_G_ values of 14,026, 18,249, and 20,508 bp respectively, from the centrophobic Alesia elements with a mean d_G_ of 1,870 and the other lineages whose d_G_ range from 4,687 (Ikeros) to 10,277 (Angela). Recombination rate correlates negatively with d_G_ (r = −0.15, P<0.001) and distinguishes the same lineages: the genomic neighbourhood of Alesia elements has a median recombination rate of 7.6 cM/Mb compared to 1.1 for RLG_BdisC152 and 3.8 for Angela. The number of segregating sites in the 10 kb flanking region, finally, also captures this fundamental distinction between LTR-RTs located mainly in low-recombination, gene-poor regions and those also occurring in more frequently recombining, gene-dense regions: the flanking regions of the former, i.e. CRM, RLG_BdisC152, Retand C, harbour less single nucleotide polymorphisms than those of more dispersed lineages like Angela, Alesia, or SIRE (Fig. 4b).

### TE lineages show great differences in GC content

A possible reason why methylation is a good predictor of the different TE lineages is that it reflects properties of the TE lineages themselves rather than being part of the ‘external’ niche of the TEs. The 32 LTR-RT lineages differ substantially in their GC content, which in turn is associated to CHG (r=0.72, P<0.001) and CpG (r=0.57, P<0.001) methylation levels. The Retand elements are particularly interesting in this regard: compared to the genome-wide median GC content of 45.6 %, the elements of the three Retand clades have GC contents of 50.3, 52.4, and 60.2 %, respectively (Table 1, Fig. 5a), and differ accordingly in methylation levels (Fig. 4b). At the other end of the spectrum, Bianca and SIRE elements have GC contents way beyond genome-wide equilibrium level: 38 and 39% respectively.

**Fig. 5:**
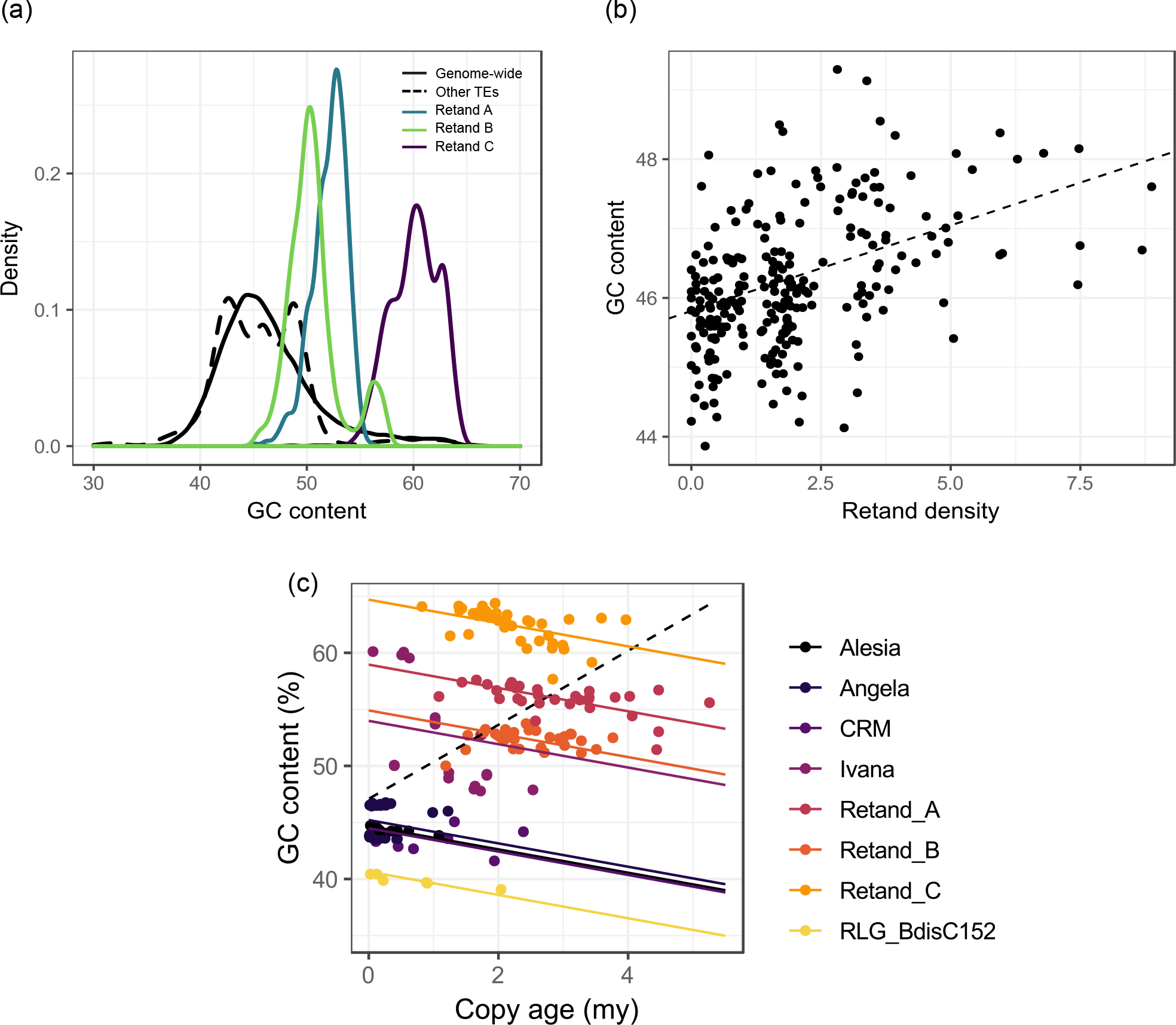
LTR-RTs and GC content. a) GC content of Retand elements compared to other LTR-RTs and genome-wide GC content in in 1 Mb bins. b) Association between GC content and the density of Retand copies in 1 Mb bins across the genome. c) GC content versus copy age for full-length elements for eight abundant lineages. The dashed line shows the slope of a simple linear model, the solid lines of the mixed effects model with the lineage as random effect.

Because Retand elements are the most abundant TEs in *B. distachyon*, their high GC content explains GC-bias on a genome-wide scale: the density of Retand elements explains more than a fifth of the variation in GC content in 1 Mb genomic windows (linear regression, r^2^=0.21, P<0.001, Fig. 5b). Recombination rate, on the other hand, which explains much of the GC bias in animal genomes (Duret & Galtier, 2009), is a much poorer predictor and has a negative effect size (linear regression, r^2^=0.08, P<0.001), which might be due to the negative correlation between Retand elements and recombination rate (r = −0.60, P < 0.001).

The high GC content of these lineages can either be an intrinsic property of the TEs or might be due to GC-biased gene conversion (gBGC) during ectopic recombination between the numerous homologous copies (Kejnovsky *et al.*, 2007). To test which explanation is more compatible with our data, we looked at the association between GC content and copy age, reflecting that gBGC would cause copies to become more GC-rich with time, while the opposite is expected when initially GC-rich copies evolve towards equilibrium GC values under the general transition/transversion bias driven by the deamination of methylated cytosines (Ossowski *et al.*, 2010). Indeed we find evidence for the prevalence of the second process, as within each lineage GC content declines with copy age at a rate of −1.03 percent per million years (standard error = 0.20, Fig. 5c).

## Discussion

In this study we investigated the LTR retrotransposon community in the Mediterranean grass *Brachypodium distachyon* in order to understand how individual TE lineages contribute to genome evolution in this species. The cast of LTR retrotransposons in *Brachypodium distachyon* includes seven major *Copia* lineages and four major *Gypsy* lineages (Fig. 1, Table 1). By comparing the properties of these lineages, we identified three ‘foundation species’ in the *B. distachyon* genome with profound effects on genome composition and dynamics: three active Angela families with an extremely fast turnover, making them an important source of within-species polymorphisms; an active non-autonomous centromeric TE family, testifying to the dynamic nature of the *B. distachyon* centromeres; and finally the abundant, old, and heavily methylated Retand copies whose presence explains much of the GC bias across the genome.

### Live fast, die young: the high-turnover Angela families

As evident in their high copy number, young age, and short half-lifes, the most active component of the *B. distachyon* genome are the three *Copia* families belonging to the Angela lineage: RLC_BdisC022, RLC_BdisC030, and RLC_BdisC024. This is the same lineage which has proliferated massively in other pooids, notably in the huge 16 Gb wheat and 5.1 Gb barley genomes. The Angela family itself was first described in wheat, where together with the closely related WIS family it is the most abundant TE in the genome (Wicker *et al.*, 2018). Similarly, the Angela family BARE1 makes up 14 percent of the diploid barley genome (Wicker *et al.*, 2017). In contrast to wheat and barley, where old intact copies are common and tend to cluster in large heterochromatic regions, the majority of Angela copies in *B. distachyon* are dispersed solo LTRs, while old intact copies virtually absent (Fig. 2).

This high number of young solo LTRs, many of which are polymorphic between Bd21 and BdTR7a, indicates that solo LTR formation is frequent in Angela families and can occur shortly after the insertion of the copy. Our comparison of solo-to-intact ratios with LTR-RT structures suggests that fast solo LTR formation in Angela families is helped by long LTRs and a comparatively short internal sequence separating them, which might facilitate ectopic within-element recombination. Such a mechanism has been suggested in a survey of LTR-RTs in eight angiosperm species (El Baidouri & Panaud, 2013). In *B. distachyon*, it would not only explain the lack of old full-length Angela copies, but also the abundance of old full-length Retand copies, since these elements have much longer internal sequences and shorter LTRs (Table 1).

More generally, and similarly to what was suggested in rice (Vitte & Panaud, 2003; Tian *et al.*, 2009), it appears that the genome of *B. distachyon* has remained small not because its TEs are idle, but because the rapid removal of its most active elements prevents a build-up of TE islands which would provide a ‘safe haven’ for TE insertion (Werren, 2011).

### Rare TEs in the genome ecosystem

An important role of LTR length in LTR-RT life history is not only suggested by high-turnover families with their long LTRs, but also by their counterparts in the community: the rare, inconspicuous TEs in the genome ecosystem. Within both superfamilies, the rarest TEs (RLG_BdisC078, Ale, Bianca, Ivana) have the shortest LTRs. The observation that all these families have few or no solo LTRs indicates that the survival of these copies in the genome might be favored by small LTRs with a low tendency to solo LTR formation.

Lineages which are rare in *B. distachyon* are also rare in other species, suggesting that their low abundance is not due to an accidental failure to proliferate, but might be the outcome of an evolutionary strategy based on inconspicuousness rather than aggressive proliferation. Also in rice and soybean, Ivana is a diverse clade with low copy numbers (Du *et al.*, 2010); the Ale lineage has been noted for its tendency to evolve a wide variety of low-copy families, while Bianca was found to be present as a low-copy single family in multiple species (Wicker & Keller, 2007). How these rare lineages manage to persist in the face of the various mutational hazards in the genome is an intriguing question. The answers to it might well modify the current picture of TE evolution, which is largely based on a few highly prolific lineages (e.g. Hawkins et al., 2006; Piegu et al., 2006).

### TEs as a source of genetic variation: burst or business as usual?

The survival curves for the three Angela families are steep (Fig. 3): half lifes are as low as 66 ky, to our knowledge the fastest rate described so far among plant TEs. Because other causes of TE ‘death’ such as purifying selection are not considered and solo LTR ages are upper-limit estimates (see Material & Methods), actual death rates might be even higher. Previous estimates at the superfamily level found half lifes of 1,265 ky for *Gypsy* and 859 ky for *Copia* elements in *B. distachyon* (International Brachypodium Initiative, 2010). Our analysis shows that these numbers misrepresent the turnover of highly active families by an order of magnitude because they average over active and inactive lineages. The same might be true for half-life estimates in other species, notably the *Copia* half-life estimates of 790 ky in rice (Wicker & Keller, 2007), 472 ky in *Arabidopsis* (Pereira, 2004), and 260 ky in *Medicago trunculata* (Wang & Liu, 2008).

Beyond providing an explanation for the small genome size of *B. distachyon*, the high sequence turnover driven by solo LTR formation advises caution in interpreting skewed age distributions as evidence for a recent activation or ‘burst’ of transposition. A left-biased age distribution is compatible with both an increased recent activity and a constant transposition rate combined with rapid removal (Dai *et al.*, 2018). Indeed, patterns of TE polymorphisms in 53 diverse natural accessions of *B. distachyon* are difficult to reconcile with transpositional bursts (Stritt *et al.*, 2018). In this previous study we found 3,627 non-reference TE insertions, mainly Angela, SIRE and RLG_BdisC152 elements segregating at low population frequencies, but no evidence for lineage-specific amplifications of single families, as might be expected if local stress had activated specific TEs (Makarevitch et al. 2015).

Because population genetic surveys of TE polymorphisms rely on discordant read-pair and splitread methods (Ewing, 2015), they treat TE insertions as presence-absence data and cannot distinguish between intact and truncated copies. By comparing the genomes of Bd21 and BdTR7a, we found that three quarters of the LTR-RTs which are not conserved between Bd21 and BdTR7a are solo LTRs, most of them belonging to the Angela lineage. This further illustrates the rapidity of solo LTR formation and suggests that a large proportion of the polymorphisms previously identified are also solo LTRs.

Finally, we observed that non-conserved TEs occur closer to genes than conserved insertions. This is consistent with purifying selection shaping the distribution of TEs in the genome, a process important in small, gene-dense genomes where the insertion of large TEs is likely to disrupt coding or regulatory sequences (Stritt *et al.*, 2018). In this light, conserved, old TEs are conserved and old because they occupy places in the genome where they interfere little with host fitness, while non-conserved TEs are primarily recent, low-frequency variants which are less likely to reach high frequencies the closer they are to genes.

### High TE activity in centromeres involves a non-autonomous family

Second to the Angela lineage, centromere-specific *Gypsy* elements are the most active part of the *B. distachyon* genome. Two families stand out with numerous young copies: RLG_BdisC039 and RLG_BdisC152. Interestingly, the latter is a non-autonomous element: it is smaller than the other CRM elements and its 3260 bp long internal sequence lacks retrotransposon ORFs as well as a chromatin targeting domain. It does contain, however, a part with high homology to the CRM families: 2000 base pairs including most of the LTR and its 3’ flanking sequence. This is the region usually containing the polymerase II promoter as well as the primer binding site for reverse transcription (Schulman, 2013). These shared regulatory motifs possibly allow the transcription and reverse transcription of RLG_BdisC152 copies. The insertion into the centromere might then be achieved by scrounging the integration complex of the autonomous CRM families.

### GC content as a distinctive feature of TE lineages

The largest contribution to genome size in *B. distachyon* comes from families of the Retand lineage. The main characteristics of these families are the high number of old full length elements, implying a low rate of solo LTR formation, and their high GC content. In Retand C, the clade with the highest copy numbers, median GC content (60.2%) is remarkably higher than the genome-wide median of 45.6 (Fig. 5). GC content in Retand copies is not only high, but also shows a phylogenetic pattern: clade A has a median content of 52.4, clade B of 50.3.

The high GC content of Retand elements has two effects. On the one hand, it implies an increased number of methylatable cytosines and higher copy methylation levels (Fig. 4b). On the other hand, because Retand elements are abundant, they alter the base composition on a genome-wide scale. Variation in GC content has been intensely studied in animals, where gBGC has emerged as the favoured explanation because it accounts for the positive association between recombination rate and GC content observed in animal genomes (Duret & Galtier, 2009). In plants, no such general pattern has emerged, possibly because the turnover of intergenic sequence is too fast for broad karyotypic patterns to emerge (Glémin *et al.*, 2014).

Transposable elements have been previously invoked to explain increased GC contents in large plant genomes, in particular of Poaceae species (Smarda *et al.*, 2014). Here we show that individual TE lineages can indeed change the GC content of a genome. More than 20% of the genome-wide variation in GC content in *B. distachyon* can be explained by the presence of Retand copies. 1.74% of the genome (4.738 Mb) were annotated as Retand copies, but because Retand elements are old and we ignored the internal sequence of truncated copies in this study, the true percentage must be much higher. As Retand elements are enriched in pericentromeric regions with low recombination rates, the presence of these elements also explains why recombination rate is negatively correlated with GC content in *B. distachyon*.

While the presence of GC-rich transposons explains particularities of the base composition and methylation landscape of *B. distachyon*, it is an open question why the base composition of these elements is so different from the rest of the genome as well as from other TE lineages. The negative association between GC content and copy age (Fig. 5c) indicates that of the two major and opposing processes affecting GC content, gBGC and the deamination of methylated cytosines (Ossowski *et al.*, 2010), the latter predominates and reduces GC content over time. It thus appears that the differing GC contents are ‘traits’ of the TE lineages themselves. An intriguing possibility is that some TE lineages have evolved elevated GC contents as a means of self-regulation (Charlesworth & Langley, 1986). The evolution of GC content as an adaptive trait would require fitness differences between TE copies differing in few base pairs. How quantitative variation in GC and methylation affects processes important for TE survival and proliferation such as transcriptional silencing, chromatin formation, and ectopic recombination rates is unclear. To clarify these issues, investigating how GC content and methylation levels vary along TE copies and how they relate to transcription factor binding sites and other functional parts of the TE could prove fruitful.

In this study we have shown that investigating LTR retrotransposon communities at the evolutionary meaningful level of lineages reveals surprising patterns which are missed when averaging over superfamilies or classes. Increasing the resolution at which TE communities are studied led us to re-appreciate the time-scale of TE dynamics: as illustrated by the Angela families, LTR-RTs can promote genome plasticity in microevolutionary time, with rapid solo LTR formation– and possibly inter-element recombination – keeping the genome small despite ongoing activity. Applying a similar approach to DNA transposons would be a logical next step to a detailed understanding of the TEs in the model grass *B. distachyon*, as *CACTA, Stowaway* and *Helitron* elements are no less diverse than LTR-RTs.

## Acknowledgements

This work was supported by the Swiss National Science Foundation (PZ00P3_154724) and the University Research Priority Programs (URPP) *Evolution in Action*. The authors would like to thank the Genetic Diversity Center at ETH Zurich for providing access to their computing resources as well as Lucy Poveda and Giancarlo Russo from the Functional Genomic Center Zurich for their support with the Bionano technology. We also thank Michael Thieme and Christian Parisod for their valuable comments on the manuscript.

## Author Contribution

CS conceived the experiment, performed the analyses and wrote the manuscript. MW performed the bisulphite sequencing and subsequent analyses. EG performed the Bionano experiment. MP assembled the genome of BdTR7a. ACR conceived and supervised the study and wrote the manuscript. All the authors read and approved the manuscript.

## Supporting Information

**Fig. S1.**
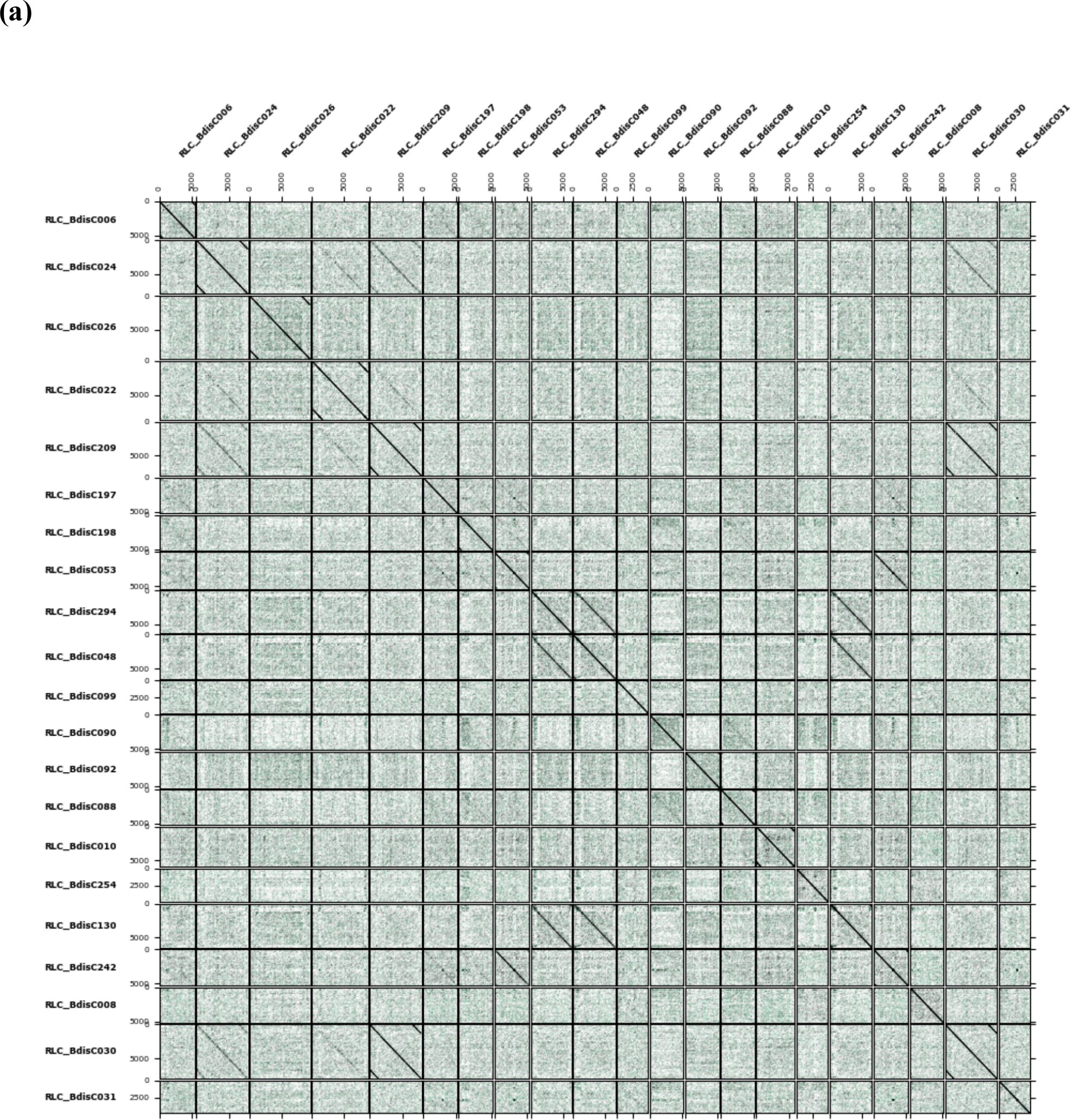
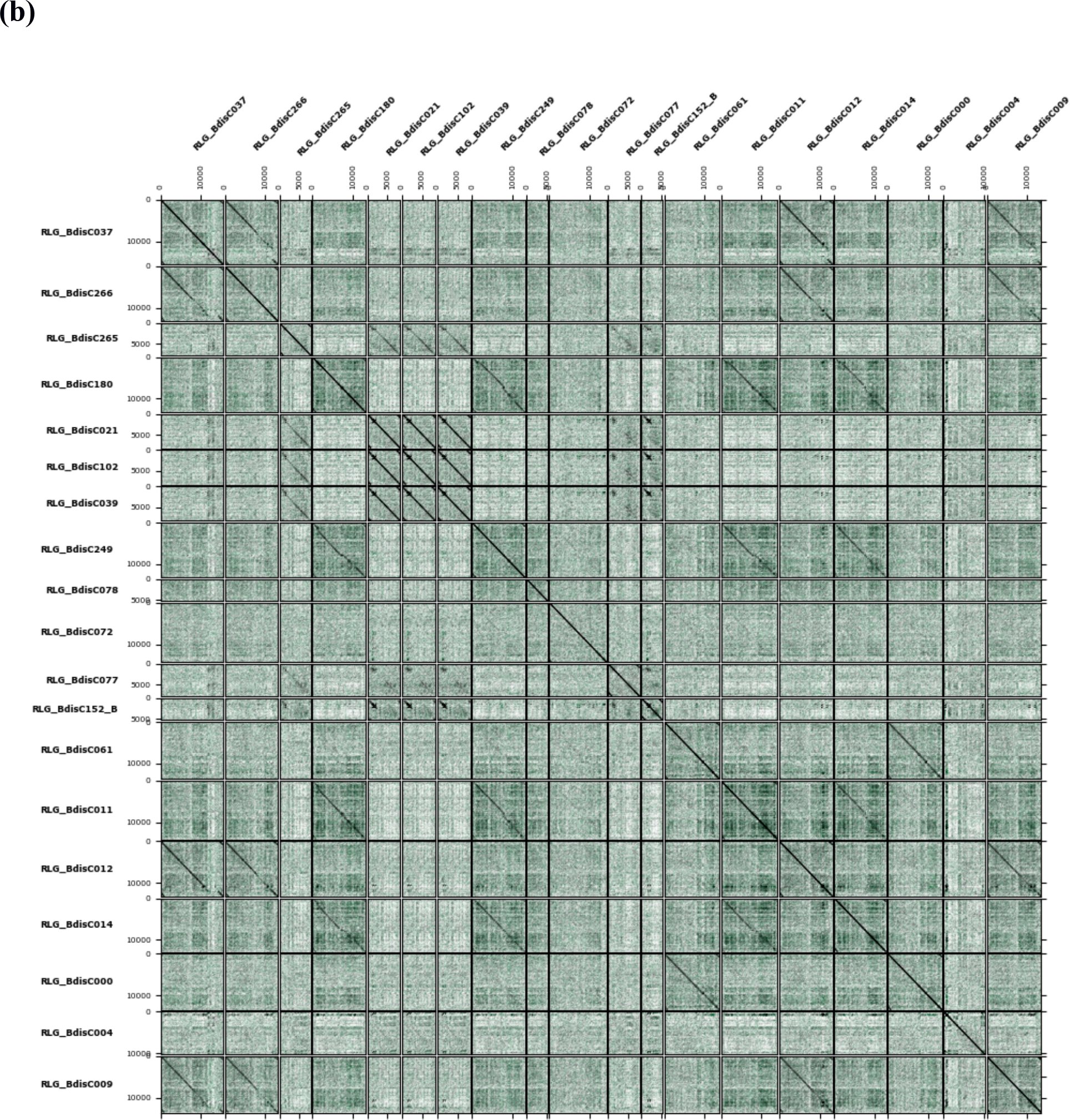
Dotplots of the 40 LTR-RT families downloaded from TREP. a) *Copia* families. The following subfamilies were merged under the name of the most abundant subfamily (underscore): RLC_BdisC030, RLC_BdisC209; RLC_BdisC053, RLC_Bdis242; RLC_Bdis294, RLC_Bdis048, RLC_Bdis130. b) For *Gypsy* familes, the following subfamilies were merged: RLG_BdisC039, RLG_BdisC102, RLG_Bdis021; RLG_BdisC000, RLG_BdisC061; RLG_BdisC266, RLG_BdisC012.

**Fig. S2.**
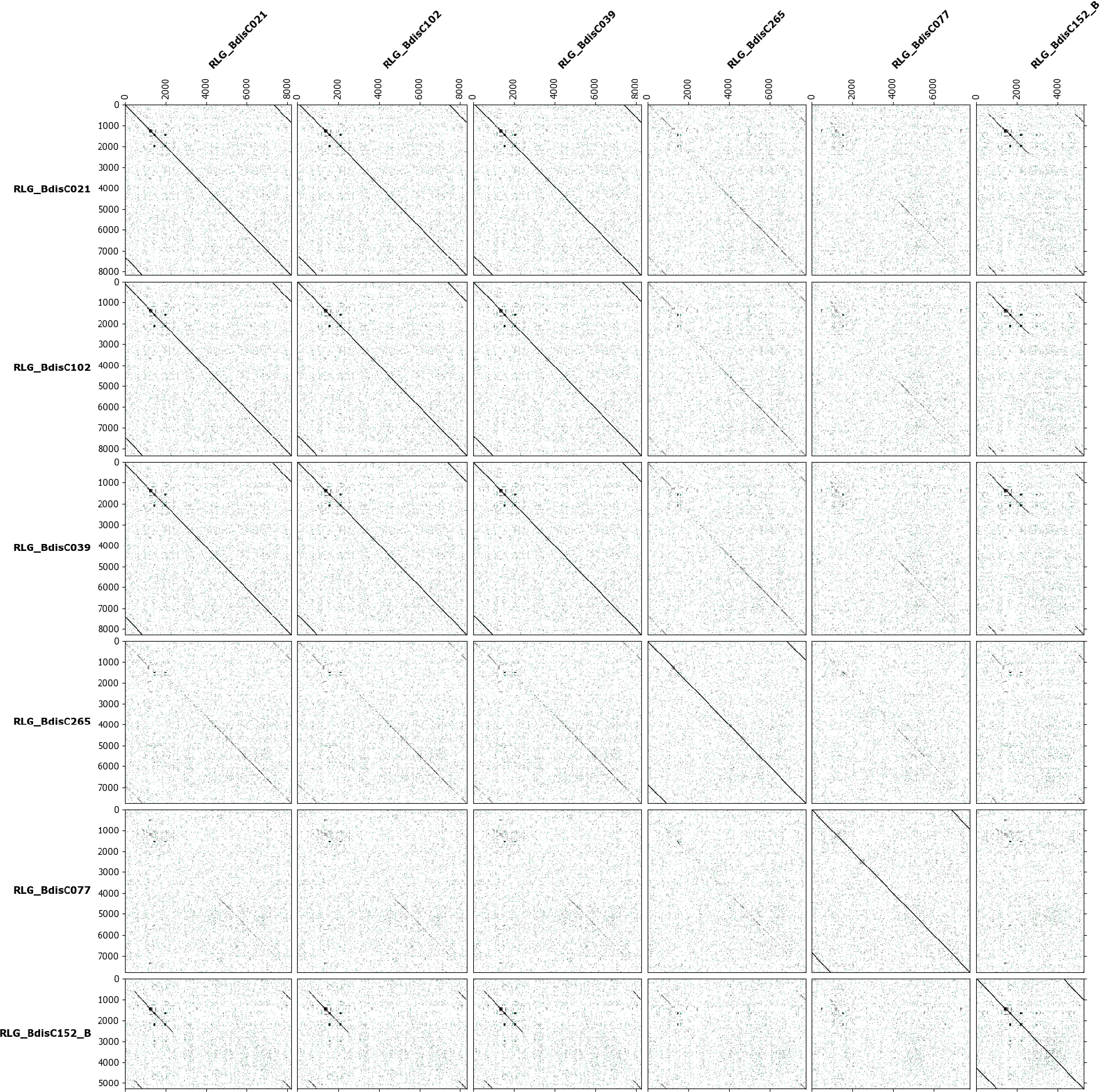
Dotplots of the consensus sequences for the five centromeric CRM families and RLG_BdisC152, which is also centromere-specific. As can be seen, the latter shares most of its LTR and part the 3’ flanking sequence with the CRM elements.

**Fig. S3.**
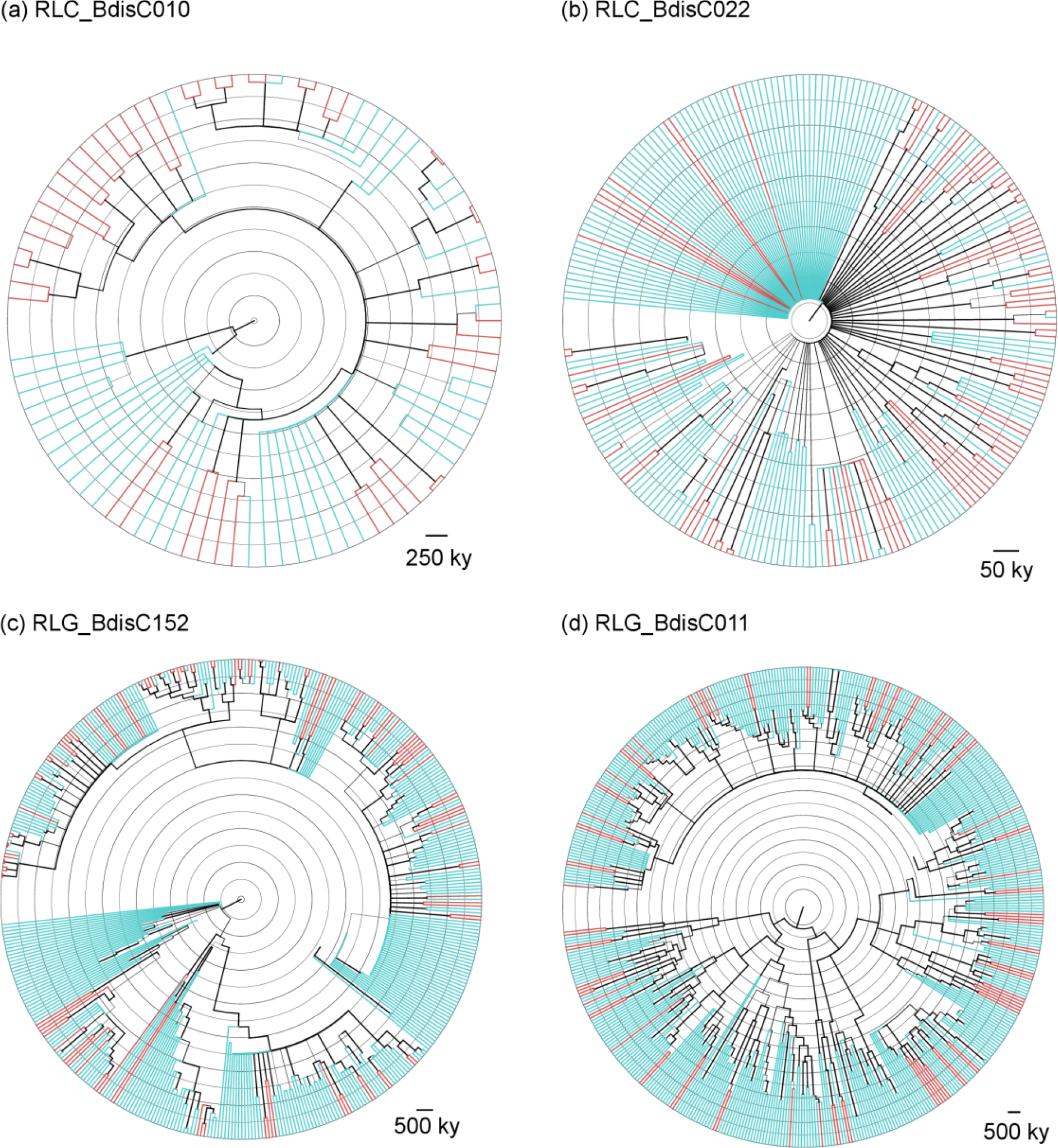
Four exemplary LTR genealogies estimated with MrBayes, showing single (blue) and paired (red) LTRs. Note the star-like phylogeny and different timescale for RLC_BdisC022, a high-turnover family. The terminal branch lengths of these trees were used as estimates for the age of the copies.

**Fig. S4.**
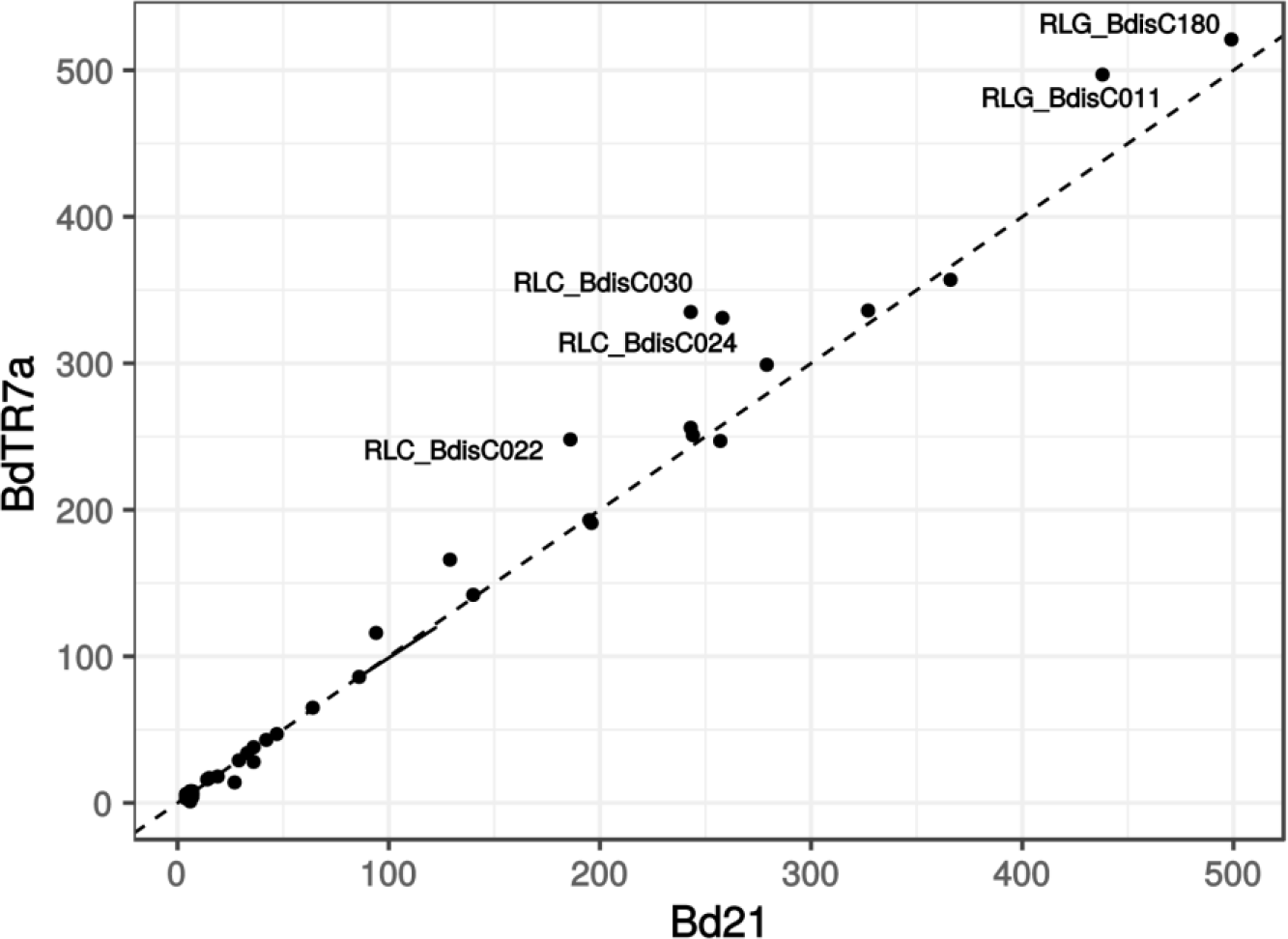
Number of annotated LTRs in Bd21 and BdTR7a. The dashed line shows the slope of 1, family names of high copy-number outliers are indicated.

**Fig. S5.**
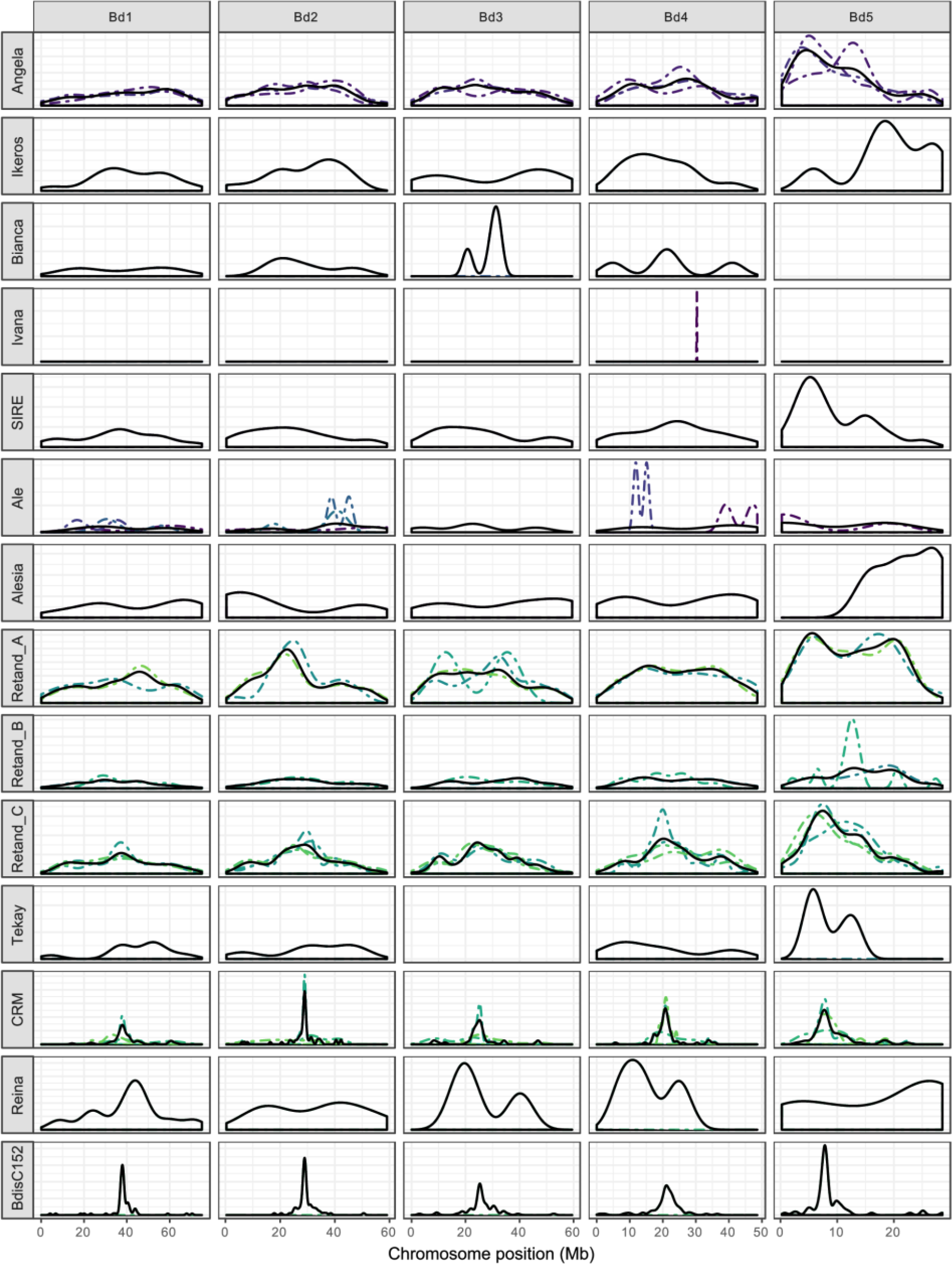
Genomic distribution of LTR retrotransposons along the five chromosomes of *B. distachyon*. Colored dashed lines show the distribution of single families with lineages, while the solid black line is the average across families.

**Fig. S6.**
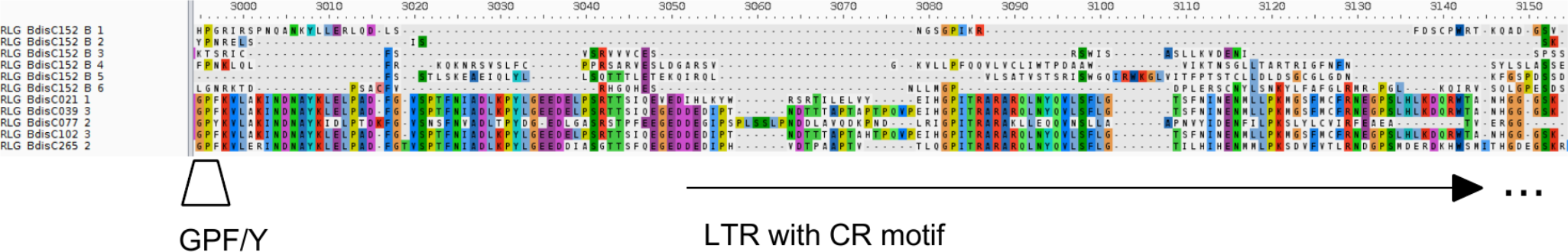
Putative chromatin targeting domain (PTD) of the CRM families of *B. distachyon*. Also shown in the alignment are the six-frame translated sequences for the centromere-specific non-autonomous family RLG_BdisC152, where the PTD is not present. See Figure 3 of Neumann et al. 2011.

**Table S1** LTR-RTs annotated in Bd21 (separate file: Bdis.LTR-RTs.Bd21.tsv)

**Table S2** LTR-RTs annotated in BdTR7a (separate file: Bdis.LTR-RTs.BdTR7a.tsv)

**Table S3.**
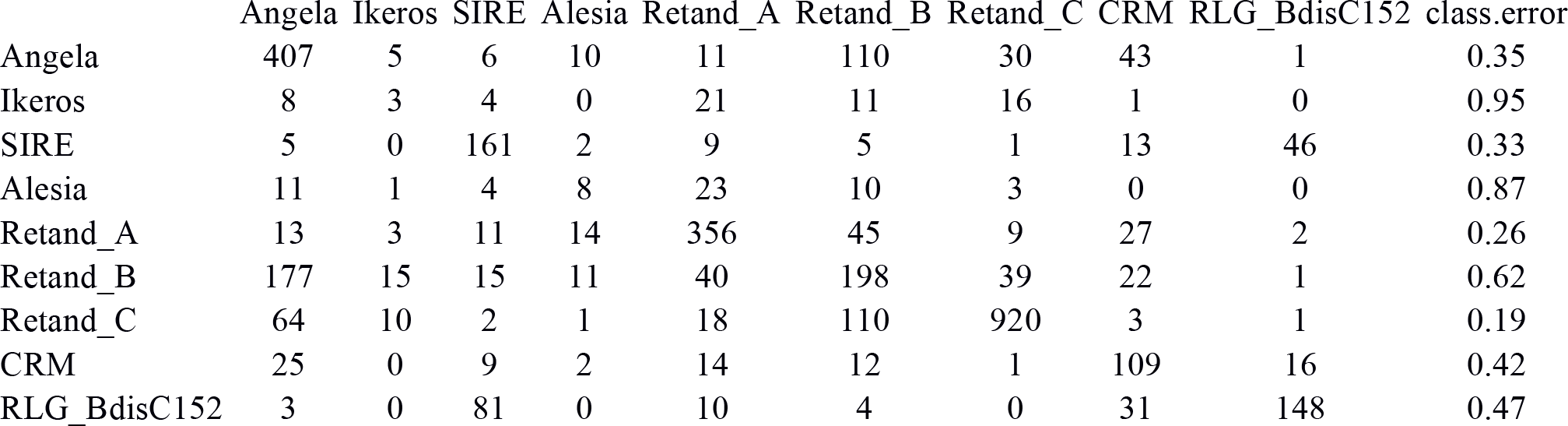
Random forest confusion matrix, showing how well the TE lineages could be distinguished from each other based on features of their genomic niche.

**Table S4.** Variable importance of the random forest model

|  | Angela | Ikeros | SIRE | Alesia | Retand_A | Retand_B | Retand_C | CRM | RLG_BdisC152 | Mean<br>Decrease<br>Accuracy |
| --- | --- | --- | --- | --- | --- | --- | --- | --- | --- | --- |
| chromosome | 0.24 | 1.77 | -0.02 | 0.42 | 0.09 | 1.00 | 1.80 | 5.06 | 5.44 | 4.26 |
| cM_per_Mb | 2.72 | 2.35 | 1.78 | 6.93 | 6.64 | 6.01 | 6.84 | 11.00 | 15.82 | 18.33 |
| dist_to_gene | 11.91 | -0.52 | 1.76 | 7.14 | 15.45 | 5.58 | 17.87 | 0.96 | 5.87 | 24.80 |
| CHG | 31.26 | 2.82 | 21.70 | 1.49 | 67.79 | 21.73 | 37.30 | 29.35 | 32.74 | 57.28 |
| CpG | 11.11 | 0.18 | 64.41 | 3.23 | 22.58 | 9.83 | 21.84 | 13.22 | 45.24 | 41.04 |
| CHH | 48.36 | 3.86 | 13.14 | 2.94 | 20.36 | 17.48 | 41.38 | 41.56 | 9.86 | 51.52 |
| S | 4.72 | 0.43 | 2.59 | 0.55 | 4.27 | 3.62 | 6.67 | 3.48 | 5.87 | 11.97 |
| $\pi$ | 6.46 | 1.54 | 3.10 | 1.13 | 3.43 | 2.84 | 6.85 | 3.81 | 5.63 | 12.38 |
| Tajima's D | 2.41 | 1.06 | 1.41 | -0.74 | 0.85 | 1.09 | 4.06 | 5.94 | 6.58 | 7.86 |
| $Z_{ns}$ | 0.48 | -1.97 | 1.23 | -1.58 | 4.60 | 4.56 | 2.47 | 3.45 | 8.43 | 8.77 |

## Methods S1: BdTR7a genome assembly

The BdTR7a assembly was created by combining PacBio sequencing with Bionano optical mapping. DNA was isolated with the Bionano Prep High Polyphenols Plant Tissue DNA Isolation Protocol (bionanogenomics.com/technology/platform-technology; August 28, 2018, date last accessed). The Bionano Sapyhr System at the Functional Genomic Center Zurich was used to produce an optical map of the genome. To do so, the extracted DNA was labelled using the PrepTM Direct Label and Stain (DLS) protocol and subsequently loaded onto a Saphyr Chip. In parallel, PacBio (Pacific Biosciences) seqencing was performed, also at the Functional Genomic Center Zurich.

Raw PacBio reads were assembled using the MARVEL assembler (Grohme *et al.*, 2018; Nowoshilow *et al.*, 2018). MARVEL consists of three major steps, namely the setup phase, patch phase and the assembly phase. In the setup phase, reads were filtered by choosing only the best read of each ZMW and requiring subsequently a minimum read length of 4 kb. The resulting 1.1 million reads (63-fold coverage) were stored in an internal database. The patch phase detects and corrects read artifacts including missed adaptors, polymerase strand jumps, chimeric reads and long low-quality read segments that are the primary impediments to long contiguous assemblies. The patched reads (55-fold coverage) were then used for the final assembly phase, which stitches short alignment artifacts resulting from bad sequencing segments within overlapping read pairs. This step was followed by repeat annotation and the generation of the overlap graph. To this end, we used the tool LAq with a quality cutoff of 27 to calculate a quality and a trim annotation track. In addition, alignments were forced through low quality regions (<200 bp) that remained in the patched reads. LArepeat in coverage auto-detection mode was used to create a repeat annotation track based on overlap coverage anomalies. The final assembled contigs were generated by touring the overlap graph. To correct base errors, we first used the correction module of MARVEL, which makes use of the final overlap graph and corrects only the reads that were used to build the contigs. Corrected contigs were further polished using PacBio’s Arrow tool (github.com/PacificBiosciences/GenomicConsensus).

Using the Bionano optical map, the 257 contigs resulting from the PacBio assembly were combined into 7 supercontigs. Finally, the these supercontigs were aligned to the Bd21 reference genome with Mummer’s nucmer algorithm (Marçais *et al.*, 2018). This step revealed that chromosomes Bd2 and Bd4 were split in two at the centromeres in the BdTR7a hybrid assembly; the final set of five chromosomes was therefore obtained by concatenating the respective supercontigs to full chromosomes.

## Methods S2: Whole-genome bisulfite sequencing

For whole-genome bisulfite sequencing, three replicates of the *B. distachyon* accession Bd21 where planted in small pots (50 ml) on a soil-sand mixture. The plants were grown for 24 days in a growth chamber with following conditions: 16 h with 200 μMol lightvolume, constant 20°C and 60 – 70% of relative air humidity.

The second and third leaf from the top were sampled, flash-frozen and disrupted using a Bead Ruptor 24 (OMNI) with following settings: T = 1:00, S = 2.50, C = 01, D = 0:00. Subsequent DNA extraction was performed with the DNeasy Plant Mini Kit (Qiagen). Around 500 ng of DNA where physically shared for 13 min with 20% of amplitude on a Q800R2 Sonicator (Qsonica). End repair and A tailing of the samples were performed with the Kapa Hyper Prep Kit (Kapa Biosystems). Methylated NimbleGen SeqCap (Roche) adapters were ligated with the above-mentioned kit and the post-ligation cleanup was performed with 0.8x AMPure XP Beads (Agencourt) and two 80% ethanol washes. Bisulfite conversion was performed with EZ DNA Methylation-Gold (ZYMO Research). The converted libraries where eluted with 20 μL for five cycles of amplification using the Kapa HiFi HotStart Uracil+ ReadyMix (Kapa Biosystems) with the Pre-LM-PCR Oligos 1 & 2 from the NimbleGen SeqCap (Roche) kit. Post-amplification cleanup was performed similarly to the above mentioned post-ligation cleanup, but with 1x of AMPure XP Beads (Agencourt). Libraries were size-selected with 0.7× of AMPure XP Beads (Agencourt) and eluted with 20 μL of 10mM Tris-HCl, pH 8.0. For library quality assessment the High Sensitivity D5000 kit on a TapeStation 2200 (Agilent Technologies) was used. For library quantification the Kapa Library Quantification Kit for Illumina Platforms (Kapa Biosystems) was used on a 7500 Fast Real-Time PCR System (Applied Biosystems). Sequencing of 151 bp paired-end reads was performed on a Illumina HiSeq 2500 (Illumina).

Reads were trimmed with trim_galore (0.4.5, bioinformatics.babraham.ac.uk/projects/trim_galore) and subsequently mapped against the reference genome of *B. distachyon* Bd21 (v.3) with Bismark (0.19.0, Krueger & Andrews, 2011) using Bowtie2 (2.3.2, Langmead & Salzberg, 2012). An average of 0.55% of the reads where identified as duplicates resulting from an overamplification of the library and removed with the deduplicate_bismark script from the Bismark. To asses the quality of the converted reads, Mbias plots (Hansen *et al.*, 2012) were produced with the bismark_methylation_extractor script using following parameters: --comprehensive -mbias_only. Visual assessment allowed to identify inconsistent sequencing of the first 2 bp and 1 bp of the 5’ end of read 1 respectively read 2. Consequently methylation levels of each covered cytosines in the specific CpG, CHG or CHH context were assessed with bismark_methylation_extractor specifying the following parameters: --comprehensive --bedGraph --CX --ignore 2 --ignore_r2 1. From reads mapping to the chloroplast sequence, we assessed an average conversion efficiency of 99%. The weighted methylation level for each annotated TE was calculated for all three context following the method described in Schultz, Schmitz & Ecker (Schultz *et al.*, 2012).

